# Genomic analysis of response to neoadjuvant chemotherapy in esophageal adenocarcinoma

**DOI:** 10.1101/2021.03.26.437144

**Authors:** Fereshteh Izadi, Benjamin P. Sharpe, Stella P. Breininger, Maria Secrier, Jane Gibson, Robert Walker, Saqib Rahman, Ginny Devonshire, Megan A Lloyd, Zoë S. Walters, Rebecca C. Fitzgerald, Matthew J. J. Rose-Zerilli, Tim J. Underwood, on behalf of OCCAMS

**Affiliations:** School of Cancer Sciences, Cancer Research UK Centre, Faculty of Medicine, University of Southampton, Southampton General Hospital, Southampton SO16 6YD, UK; UCL Genetics Institute, Division of Biosciences, University College London, Gower Street, London, WC1E 6BT, UK; Institute for Life Science, University of Southampton, SO17 1BJ, UK; Cancer Research UK Cambridge Institute, University of Cambridge, Cambridge, UK

**Keywords:** Esophageal adenocarcinoma, chemotherapy, mutation, response, genomics, *NAV3*

## Abstract

Neoadjuvant therapy followed by surgery is the standard of care for locally advanced esophageal adenocarcinoma (EAC). Unfortunately, response to neoadjuvant chemotherapy (NAC) is poor (<20%), as is the overall survival benefit at 5 years (5%). The EAC genome is complex and heterogeneous between patients, and it is not yet understood whether specific mutational patterns may result in chemotherapy sensitivity or resistance. To identify associations between genomic events and response to NAC in EAC, a comparative genomic analysis was performed in 65 patients with extensive clinical and pathological annotation using whole-genome sequencing (WGS). We defined response using Mandard Tumor Regression Grade (TRG), with responders classified as TRG1-2 (n=27) and non-responders classified as TRG4-5 (n=38). We report a higher non-synonymous mutation burden in responders (median 2.08/Mb vs 1.70/Mb, *P*=0.036) and elevated copy number variation in non-responders (282 vs 136/patient, *P*<0.001). We identified copy number variants unique to each group in our cohort, with cell cycle (*CDKN2A, CCND1*), c-Myc (*MYC*), RTK/PIK3 (*KRAS, EGFR*) and gastrointestinal differentiation (*GATA6*) pathway genes being specifically altered in non-responders. Of note, *NAV3* mutations were exclusively present in the non-responder group with a frequency of 22%. Thus, lower mutation burden, higher chromosomal instability and specific copy number alterations are associated with resistance to NAC.

## Introduction

Esophageal adenocarcinoma (EAC) is a cancer of unmet clinical need. Patients with locally advanced EAC suitable for curative treatment receive neo-adjuvant chemoradiotherapy or neo-adjuvant chemotherapy (NAC) with or without adjuvant chemotherapy as standard of care. Randomized trials of NAC have consistently shown survival benefits for patients (Ronellenfitsch et al. 2013; Girling et al. 2002; Cunningham et al. 2006; Allum et al. 2009; Al-Batran et al. 2019). However, this survival advantage (5% at 5 years) (Smyth et al. 2017) is not due to an incremental improvement in outcome for all patients, but instead driven by a very good response in less than 20% of patients (Noble et al. 2017; Sjoquist et al. 2011). Primary tumor regression following neoadjuvant therapy (NAT) can be measured using the Mandard Tumor Regression Grade (TRG) in resected specimens after surgery (Mandard et al. 1994; Tan et al. 2016; Tao et al. 2015) and is informative for both disease-free and overall survival (Wong and Law 2017; Tan et al. 2016; Tao et al. 2015). Genetic mechanisms associated with tumor response to NAT have been assessed in a variety of different cancer types (Chakiba et al. 2014; Cramer et al. 2018; Greenbaum et al. 2019; Höglander et al. 2018; Lesurf et al. 2017; Li et al. 2020; Zhu et al. 2020) including rectal adenocarcinoma, but have not been widely investigated in EAC. Predictive biomarkers of response following NAT have been proposed for EAC, including functional imaging and expression of genes regulating apoptosis, angiogenesis, cell cycle, and DNA repair as well as growth factors and their receptors, but none have approached clinical practice (Tan et al. 2016; Tao et al. 2015).

EAC genomes are characterized by a high degree of chromosomal instability (Nones et al. 2014; Secrier et al. 2016), and large-scale genomic studies, such as those conducted by the OCCAMs UK consortium using whole genome sequencing to contribute the International Cancer Genome effort, have identified key driver genes and clinically relevant biomarkers for prognostication (Secrier et al. 2016; Frankell et al. 2019). This large cohort with extensive data on treatment and clinic-pathological response provides an ideal opportunity to investigate the predictors of response and resistance to chemotherapy. To date, only two studies using whole exome sequencing have investigated genetic features associated with response to NAC in which genetic bottlenecks, intratumor heterogeneity and early chromosomal instability were found to be related to NAC response/resistance in EAC (Findlay et al. 2016; Murugaesu et al. 2015). These studies provide key insights into the genomic evolution of EAC through NAC and the changes in the genome architecture following clinical response. There is a need for studies to further characterize the whole genomic landscape in EAC at the time of diagnosis (pre-treatment) to enable identification of predictive biomarkers for response to NAC and to identify the consequences of genomic lesions suitable for novel interventions.

Here, we describe results from whole genome sequencing (WGS) of pre-treatment biopsies from 65 EAC patients treated with NAC and surgery, alongside RNA-seq, to investigate the genetic features associated with NAC response. We describe a hierarchical approach to the comparative analysis of the genomes of EAC responders and non-responders, starting with total mutational burden, continuing through large-scale chromosomal events and on to driver gene mutations, before defining the key genomic differences between groups and their potential for possible therapeutic intervention.

## Results

### Patient characteristics and overall study design

In total, 65 cases from the well-curated OCCAMS consortium multi-center dataset (Secrier et al. 2016; Frankell et al. 2019) were classified into two groups based on Mandard Tumor Regression Grading (TRG): 27 responders (TRG1 (n=18) and TRG2 (n=9)) and 38 non-responders (TRG4 (n=28) and TRG5 (n=10)) (**Fig. 1A**). We excluded TRG3 classified cases because of their prognostically heterogeneous behaviour (Mancini et al. 2018). A summary of the clinicopathological data for the cohort is shown in **Table 1** with full details available in (**Table S1**). Median follow-up in the cohort was 56.7 months (1.5 - 78.8 months). In line with our previous multi-centre cohort study (Noble et al. 2017), TRG defined responders had favourable prognosis compared to non-responders with a significantly longer overall survival (78.5 vs. 33.8 months, *P* < 0.001, **Fig. 1B**). As expected, following NAC, non-responders had higher pathological TNM stage (ypT and ypN) compared to responders (χ^2^ test, *P* = 0.001, **Table 1**) but there was no difference in pre-NAC TNM stage (Rice et al. 2017). All 65 patients had WGS data generated from endoscopic biopsies and matched germline DNA taken at the time of cancer diagnosis and before any treatment. In total, 9 responders and 21 non-responders had matched RNA-seq data to complement the WGS dataset according to the availability of tissue.

**Figure 1.**
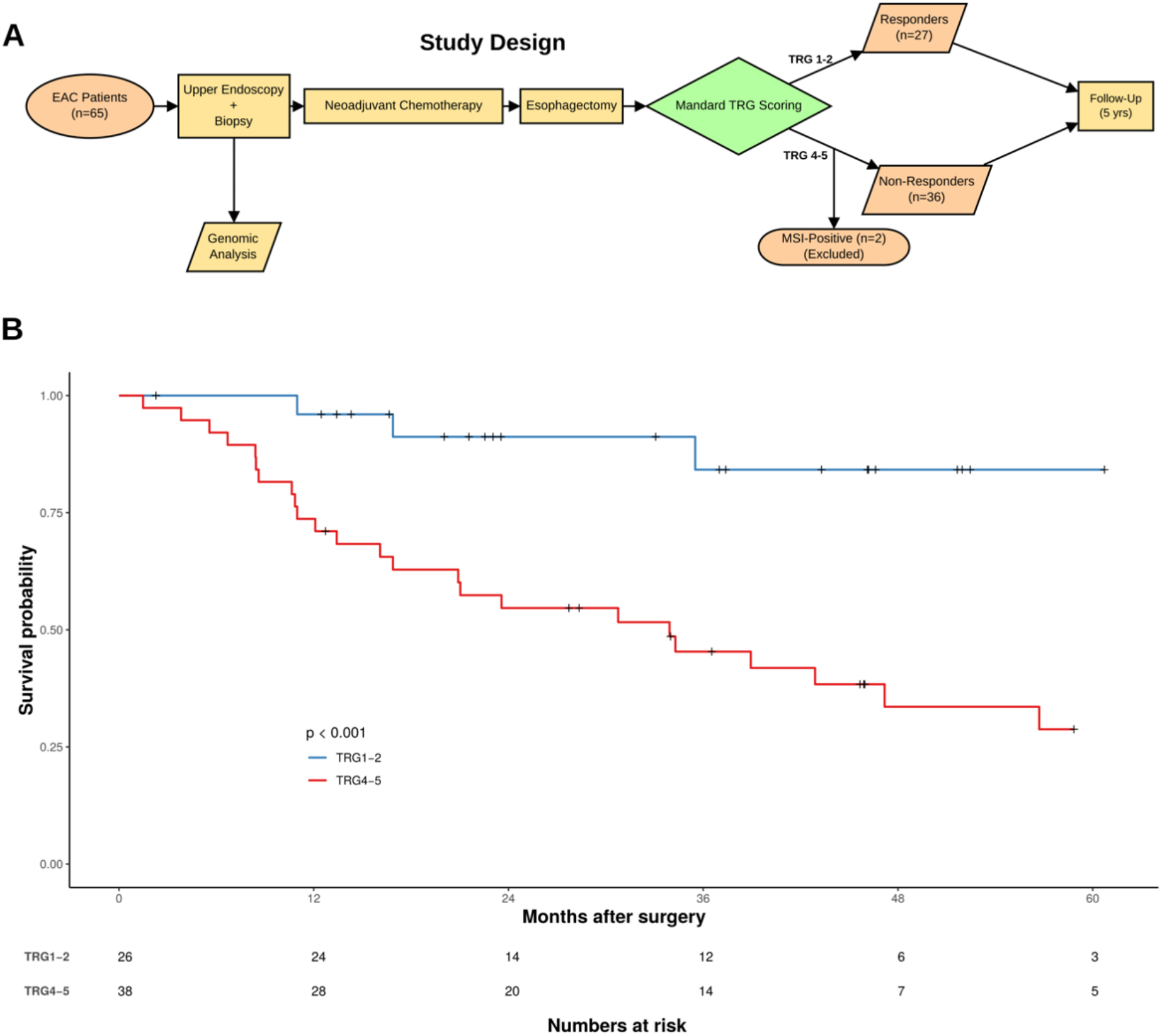
Outline of the cohort and analyses performed. (**A**) Description of the study design. (**B**) Kaplan–Meier of overall survival (n=64) for responders (blue line) and non-responders (red line). Number of cases at risk are detailed in the table.

**Table 1:** Clinicopathological data for the study cohort according to response to NAC.

| Variable | Category | Non-responders (n=38) | Responders (n=27) | Overall (n=65) | P (X<sup>2</sup> test) |
| --- | --- | --- | --- | --- | --- |
| **Age** |  | 66.25 | 64.30 | 65.00 | 0.739¶ |
| **Gender** | Female | 5 (13.2) | 2 (7.4) | 7 (10.8) | 0.741 |
|  | Male | 33 (86.8) | 25 (92.6) | 58 (89.2) |  |
| **cT Stage** | T1 | 0 (0.0) | 1 (3.7) | 1 (1.5) | 0.485 |
|  | T2 | 5 (13.2) | 5 (18.5) | 10 (15.4) |  |
|  | T3 | 31 (81.6) | 19 (70.4) | 50 (76.9) |  |
|  | T4 | 2 (5.3) | 1 (3.7) | 3 (4.6) |  |
|  | Missing | 0 (0.0) | 1 (3.7) | 1 (1.5) |  |
| **cN Stage** | N0 | 8 (21.1) | 7 (25.9) | 15 (23.1) | 0.494 |
|  | N1 | 24 (63.2) | 12 (44.4) | 36 (55.4) |  |
|  | N2 | 5 (13.2) | 6 (22.2) | 11 (16.9) |  |
|  | N3 | 1 (2.6) | 1 (3.7) | 2 (3.1) |  |
|  | Missing | 0 (0.0) | 1 (3.7) | 1 (1.5) |  |
| **Tumor Location** | GOJ | 19 (50.0) | 16 (59.3) | 35 (53.8) | 0.627 |
|  | Esophagus | 19 (50.0) | 11 (40.7) | 30 (46.2) |  |
| **ypT Stage** | T0 | 1 (2.6) | 17 (63.0) | 18 (27.7) | <0.001* |
|  | T1 | 3 (7.9) | 5 (18.5) | 8 (12.3) |  |
|  | T2 | 4 (10.5) | 1 (3.7) | 5 (7.7) |  |
|  | T3 | 24 (63.2) | 4 (14.8) | 28 (43.1) |  |
|  | T4 | 6 (15.8) | 0 (0.0) | 6 (9.2) |  |
| **ypN Stage** | N0 | 9 (23.7) | 17 (63.0) | 26 (40.0) | <0.001* |
|  | N1 | 6 (15.8) | 4 (14.8) | 10 (15.4) |  |
|  | N2 | 14 (36.8) | 2 (7.4) | 16 (24.6) |  |
|  | N3 | 8 (21.1) | 0 (0.0) | 8 (12.3) |  |
|  | Missing | 1 (2.6) | 4 (14.8) | 5 (7.7) |  |
| **ypM Stage** | M0 | 35 (92.1) | 27 (100.0) | 62 (95.4) | 0.371 |
|  | M1 | 3 (7.9) | 0 (0.0) | 3 (4.6) |  |

Data presented as absolute number (%) and median (IQR), \*P<0.05. ¶ indicates a Mann–Whitney U test p-value.

### Mutational profiles associated with response to NAC

In order to establish an overview of the mutational landscape associated with response to NAC, we compared the genomes of responders and non-responders. We performed assessment of somatic copy number variations (CNVs), small insertions/deletions (indels) and variant calling of somatic single nucleotide variants (SNVs) as previously described (Frankell et al. 2019; Li et al. 2018; Noorani et al. 2017; Secrier et al. 2016). Two non-responders were found to have microsatellite instability (MSI) (Patient 27, score= 5.63 and Patient 29, score= 3.44) and these were excluded because MSI-High tumors are known to respond to immune checkpoint blockade, providing a potential treatment pathway for these patients (van Velzen et al. 2020).

Initially, we investigated overall mutation burden in responders and non-responders on a genome-wide level. We identified a median of 104 (3-286) non-synonymous mutations per tumor genome in responders, compared to 85 (1-171) mutations in non-responders and the mutation frequency per megabase (Mb) was higher in the responder group (2.08 (range: 0.14–3.66) vs 1.7 (range: 0.02–3.42); Fig. 2A, Wilcoxon rank sum test, *P* = 0.036).

**Figure 2.**
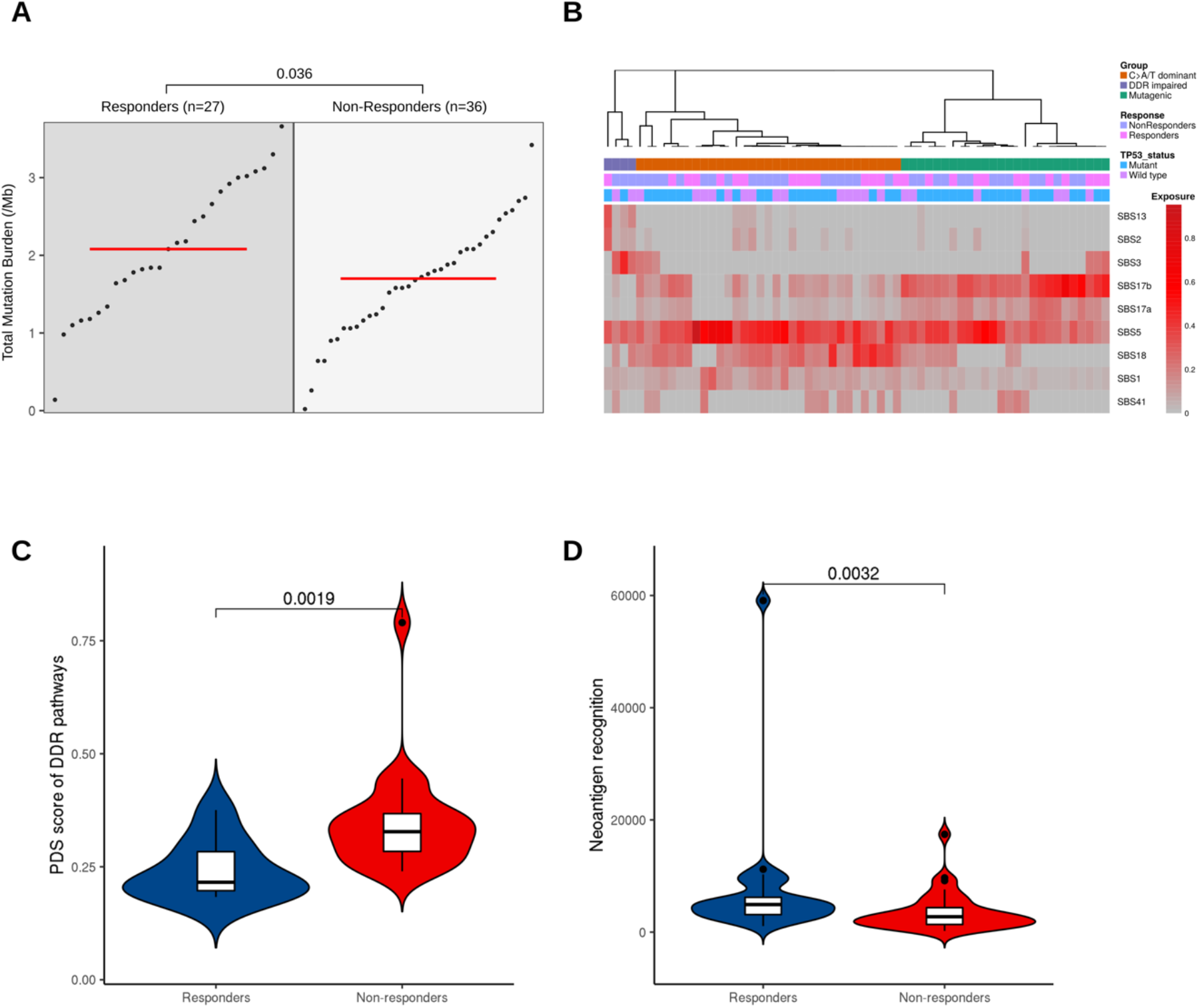
Mutational landscape of NAC response. Responders have a higher mutation burden and neoantigen recognition potential. (**A**) Group dot plot of mutation per megabase. Each dot represents a patient with the red line marking the group median. (**B**) Clustering of the 9 mutational signatures in our patient samples as previously described by Secrier et al (2016). (**C**) Pathway Deregulation Scores (PDS) calculated using gene expression values (log2 normalized) of available RNA-seq samples for DDR pathways. (**D**) Neoantigen recognition potential scores. Only neoantigens with recognition potential above one are shown.

To investigate the mutational profile of SNVs in the trinucleotide context in our cohort, we performed mutational signature extraction using SigProfiler (Alexandrov et al. 2020). Nine mutational signatures were defined in our cohort (Fig. 2B). We hierarchically clustered our cases and in agreement with prior studies (Secrier et al. 2016), three main subgroups were observed corresponding to predominant signatures: these are classified as C>A/T dominant (SBS1/5 and SBS18); DDR impaired (SBS3); and mutagenic (SBS17A/B) subgroups. There was no significant difference in the proportion of responders and non-responders between subgroups (Chi square test, *P* = 0.4). However, the DDR impaired subgroup is defined by signature 3 mutations, and the majority of these tumors (3/4) were nonresponders. Signature 3 is associated with failure of DNA double-strand break-repair by homologous recombination, which could lead to chromosomal instabilities. Consequently, we assessed the dysregulation of DDR pathways using gene expression data of available RNA-seq samples (n=9 responders, n=21 non-responders) and created a pathways dysregulation score (PDS) using Pathifier (Drier et al. 2013). Non-responders exhibit greater dysregulation in DDR pathways compared to responders (Fig. 2C, Wilcoxon rank sum test, *P* = 0.002).

Having observed higher mutational burden in responders, we hypothesised that this would also be correlated with a greater neoantigen load. We used NeoPredPipe (Schenck et al. 2019), a predictive tool, to identify tumor neoantigens using binding affinity for patient-specific class I human lymphocyte antigen (HLA) alleles. We next calculated the neoantigen recognition potential by quantifying the peptides that displayed high affinity binding in tumors, but had no predicted binding in the matched normal sample (Luksza et al. 2017). Considering only those samples with recognition potential value above 1, responders had a significantly higher neoantigen recognition potential score (Wilcoxon ranksum test, *P* < 0.001, Fig. 2D), possibly supporting previous associations between CD8+ tumor infiltrating lymphocyte levels and improved survival in EAC following NAC (Noble et al. 2016; Secrier et al. 2016).

### Non-responders have more chromosomal instability and unique copy number alterations

We next moved to consider chromosomal and copy number events and correlate these with mRNA expression, where possible, before considering point mutations. To investigate correlations between chromosomal instability (CIN) and response to NAC, we measured the proportion of the genome affected by copy number alteration. Non-responders exhibited a higher level of CIN as evidenced by a higher Genomic Instability Index (GII) (Do Canto et al. 2019) (Wilcoxon rank sum test, *P* < 0.001, Fig. 3A). To confirm these findings, we evaluated the CIN70 signature in matched RNA-Seq data, a gene signature whose expression was consistently correlated with total functional aneuploidy across multiple cancer types (Carter et al. 2006). We found a higher CIN70 signature in non-responders, but this was not significant (*P* = 0.064), likely due to the small size of the RNA-Seq cohort.

**Figure 3.**
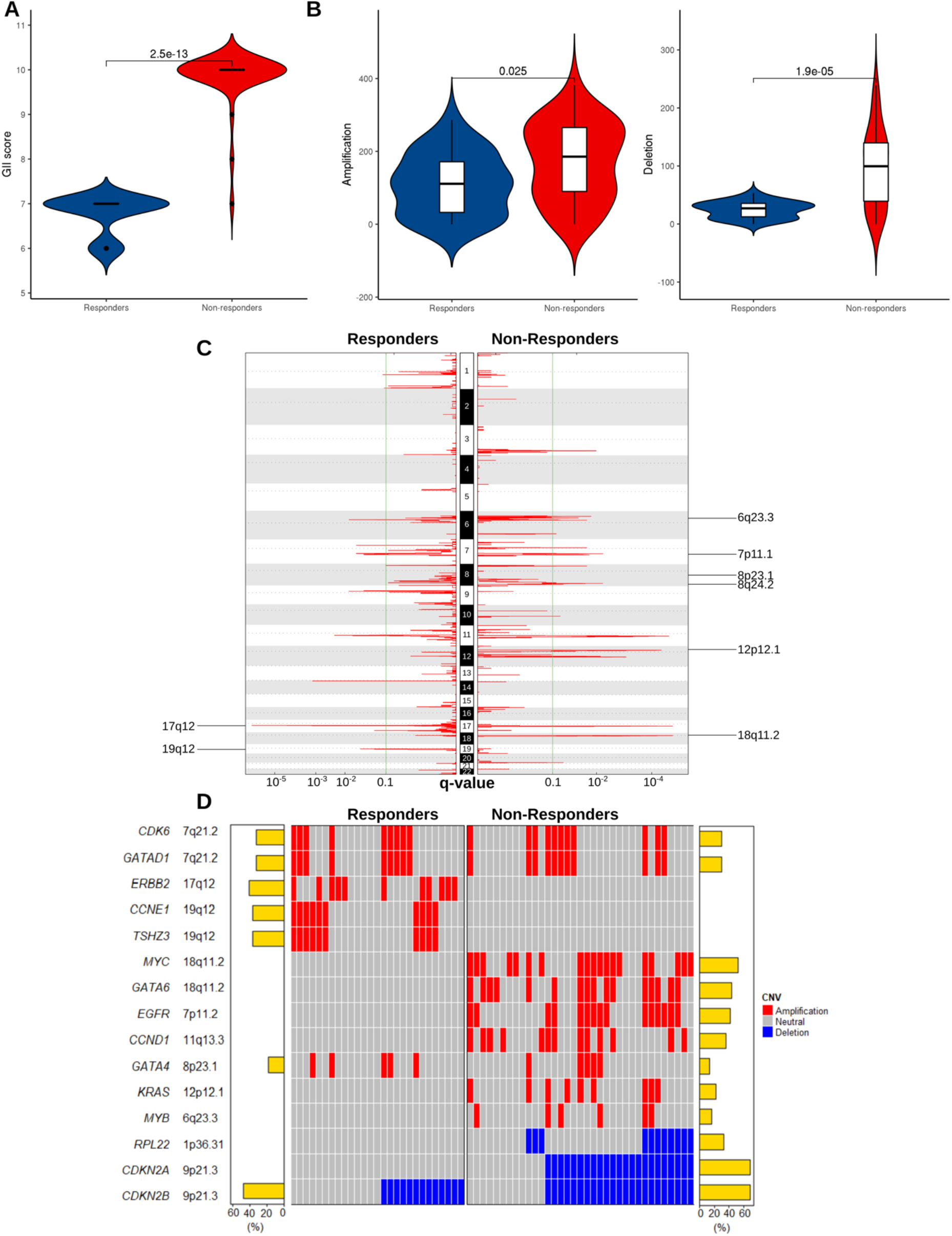
Non-responders have less stable genomes and unique patterns of copy number change in EAC driver genes. (**A**) Proportions of the genome affected by copy-number changes (Genomic Instability Index, GII). Non-responders showed a higher level of genomic instability (*P*=2.5e-13). (**B**) Violin plots depicting the frequency of all amplifications and deletions in responders and nonresponders. (**C**) Amplified peak regions across the genome plotted for responders versus nonresponders (n=63) using GISTIC2.0 (FDR < 0.1). Amplifications unique to each group are labelled. (**D**) Oncoplot of recurrently amplified/deleted EAC drivers among responders and non-responders identified by GISTIC2.0 (FDR < 0.1).

We then identified recurrently amplified or deleted regions using GISTIC2.0 (Mermel et al. 2011). In responders, a total of 3,626 CNVs were detected (median 136/patient, range: 0–292) including 2,961 amplifications (median 115/patient, range: 0–286) and 665 deletions per case (median 28/patient, range: 5–53). In non-responders, there were a total of 9,637 CNVs (median 282/patient, range: 5-504) including 6,198 amplifications (median 185.5/patient, range: 0-382) and 3,439 deletions (median 99.5/patient, range: 0-239). The total CNVs and the number of amplifications and deletions were higher in non-responders (Wilcoxon rank sum test, *P*-values < 0.001, 0.025 and 0.001 respectively, Fig. 3B). At the chromosomal arm level, we found recurrent amplifications of chr 20q in 48% of responders (FDR < 0.1), while we identified no significant amplification in non-responders (Supplemental Tables S2, S3). Furthermore, we found 13 and 20 significant deletions of chromosomal arms in responders and non-responders respectively. Deletion of 14p was unique to responders, and deletions of 8p, 9q, 9p, 10q, 15q, 16p, 19q, and 22q were unique to this cohort of non-responders, showing a higher level of large-scale deletion in non-responders.

We next looked at copy number alterations in 76 previously validated EAC driver genes (Frankell et al. 2019). We restricted our initial analysis to these 76 genes as they have been comprehensively analysed and verified in contemporaneous and clinically relevant cohorts in addition to downstream functional biological assessment. We identified significantly amplified or deleted peaks for the two groups (FDR < 0.1, Fig. 3C&D, Supplementary Tables S4-5). Distinct focal amplifications and deletions in EAC driver genes are illustrated in Figure 3D. The responders contained two unique amplification peaks: 17q12, containing *ERBB2* (FDR < 0.001) and 19q12, containing *CCNE1* and *TSHZ3* (FDR = 0.003) (Supplemental Table S4). These focal amplifications contrast with observed chromosome arm deletions in 19q observed in non-responders. Meanwhile, the non-responders contained more unique amplification peaks: 18q11.2 containing *GATA6* (FDR < 0.001); 7p11.2 containing *EGFR* (FDR = 0.018); 11q13.3.2 containing *CCND1* (FDR < 0.001); 12p12.1 containing *KRAS* (FDR < 0.001); 6q23.3 containing *MYB* (FDR = 0.085); and 8q24.21 containing *MYC* (FDR = 0.007) (Supplemental Table S5). Focal amplifications in MYC and GATA6 in non-responders contrast with our findings of arm level deletions in responders at the same chromosome arm, 18q. We investigated whether copy number changes in driver genes were co-occurrent and found that *GATA6* and *EGFR* were co-occurrent in non-responders (Fisher’s exact test, *P* = 0.002, Supplemental Tables S6-7).

### mRNA expression level supports the dysregulation of EAC driver genes in non-responders

We reasoned that if these amplification/deletion peaks played a role in affecting the response to NAC, then we would observe corresponding signals in their related downstream pathways and patient survival would be affected. To do this we used matched RNA-seq data (n=9 responders, and n=21 non-responders) and compared the FPKM values (Fragments Per Kilobase of transcript per Million mapped reads) of recurrently amplified and deleted EAC driver genes between groups. Consistent with copy number alterations, we observed the upregulation of *CDK6* (Wilcoxon rank sum test, *P* = 0.004), *CCND1* (*P* = 0.004), *GATA4* (*P* = 0.037) and *MYC* (*P* < 0.001) at the transcript level in non-responders (Fig. 4A). Furthermore, patients with amplification at the corresponding chromosomal regions of cell cycle regulators had a worse prognosis with a shorter overall survival, including *CCND1* (median survival 20.8 months in *CCND1* amplified samples versus 78.5 months in *CCND1* neutral samples, *P* = 0.007), *CDK6* (median survival 33.8 months in *CDK6* amplified samples versus 73.0 months in *CDK6* neutral samples, *P* = 0.01) and deletion of regions harbouring *CDKN2A* (median survival 33.8 months in *CDKN2A* deleted samples versus 73.0 months in *CDKN2A* neutral samples, *P* = 0.01) (Fig. 4B). Among pathways related to these genes, only *MYC* signalling was significantly enriched in the non-responders using Gene Set Enrichment Analysis (GSEA) (FDR = 0.04, Fig. 4C and Supplemental Table S8-9). Although we observed a significantly elevated expression of *MYC* in non-responders, *MYC* amplification was not significantly associated with overall survival (median survival 30.7 months in *MYC* amplified samples versus 73.0 months in *MYC* neutral samples, *P* = 0.068). Overall, we found no significant influence of these copy number changes on overall survival in responders or nonresponders alone, as our study was underpowered for these comparisons (Supplementary Figure S1). However, in responders CDK6 amplification was associated with shorter overall survival (median survival 35.4 months in CDK6 amplified samples versus 78.5 months in CDK6 neutral samples, P = 0.0011).

**Figure 4.**
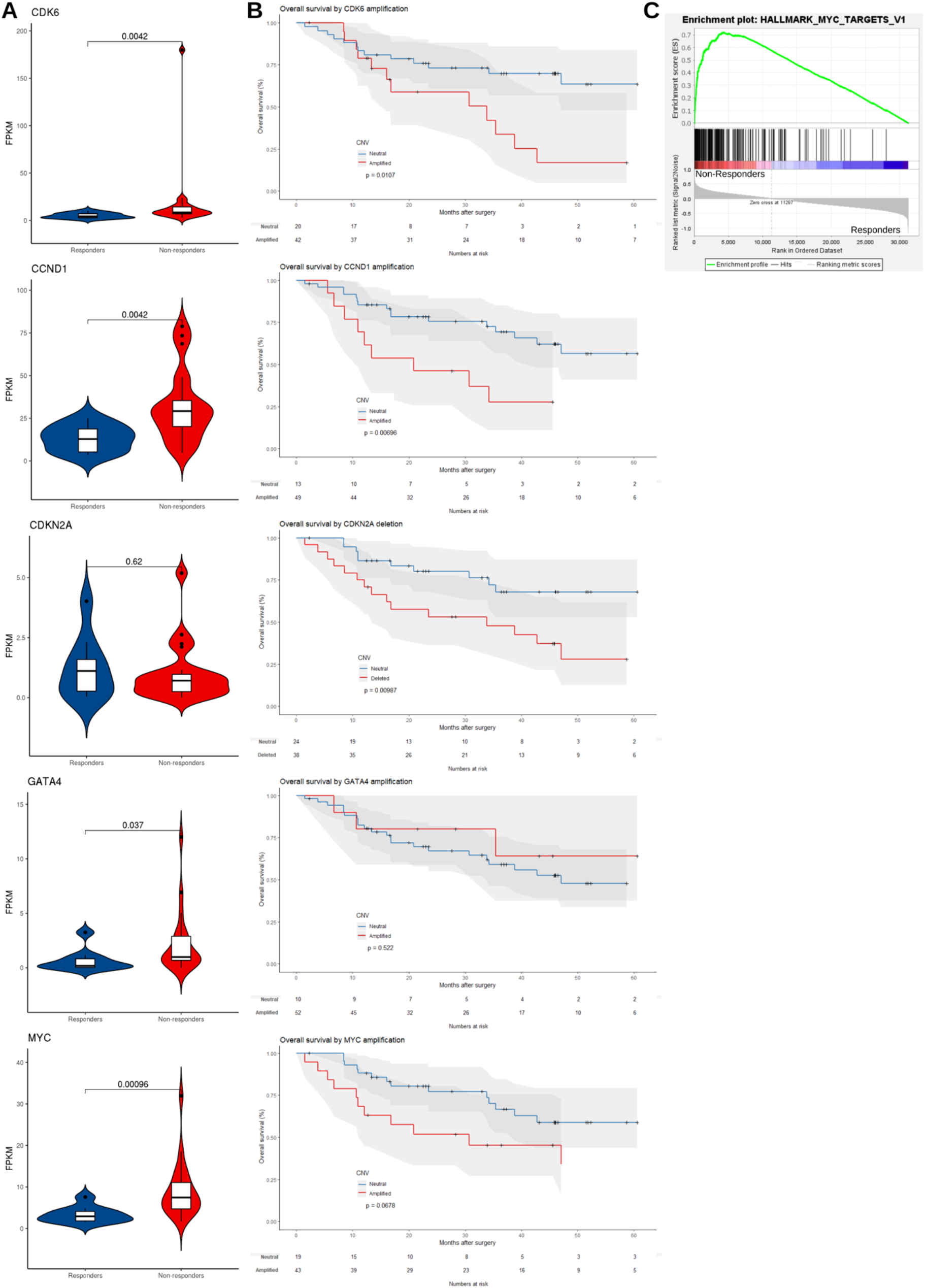
Amplified EAC driver genes are overexpressed in non-responders and copy number alterations associate with poor survival. (**A**) Violin plots comparing mRNA expression levels (FPKM) in matched RNA-Seq data (n=9 responders, n=21 non-responders) for copy number altered EAC driver genes in responders and non-responders. *P*-values were based on one-tailed Wilcoxon rank sum test. (**B**) Kaplan-Meier plot comparing overall survival of patients with *CDK6, CCND1, CDKN2A, GATA4* and *MYC* copy number changes (red) versus neutral patients (blue). (**C**) Gene Set Enrichment Analysis (GSEA) of *MYC* target genes in available RNA-Seq data (n=30).

### Mutated driver genes differ between responders and non-responders

Having established the potential importance of chromosomal level structural variation and gene level copy number variation in response to NAC in EAC, we next moved to assess SNVs, starting with known EAC driver genes (Fig. 5A). WGS data showed that 96.2% of responders and 94.3% of non-responders carried at least one non-synonymous somatic mutation in these EAC driver genes. As expected, *TP53* (65%), *CDKN2A* (16%), *SMAD4* (15%) and ARID1A (8%) were highly mutated in this cohort (Frankell et al. 2019). *NAV3* was exclusively mutated in non-responders (8/36, 22%, Fisher’s exact test, *P* = 0.01)(Fig. 5B). We used the Ensembl Variant Effect Predictor (McLaren et al. 2016) to predict mutational consequences and found that several mutations, including *NAV3* p.V142G and p.D2366N, are likely to be functionally deleterious (Supplementary Table S10). We also found mutations in *KCNQ3* (4/36 patients, 11%), *LRRK2* (4/36 patients, 11%), *KRAS* (3/36 patients, 8%), and *PBRM1* (2/36 patients, 6%) that were unique to non-responders, but these were not statistically significant.

**Figure 5.**
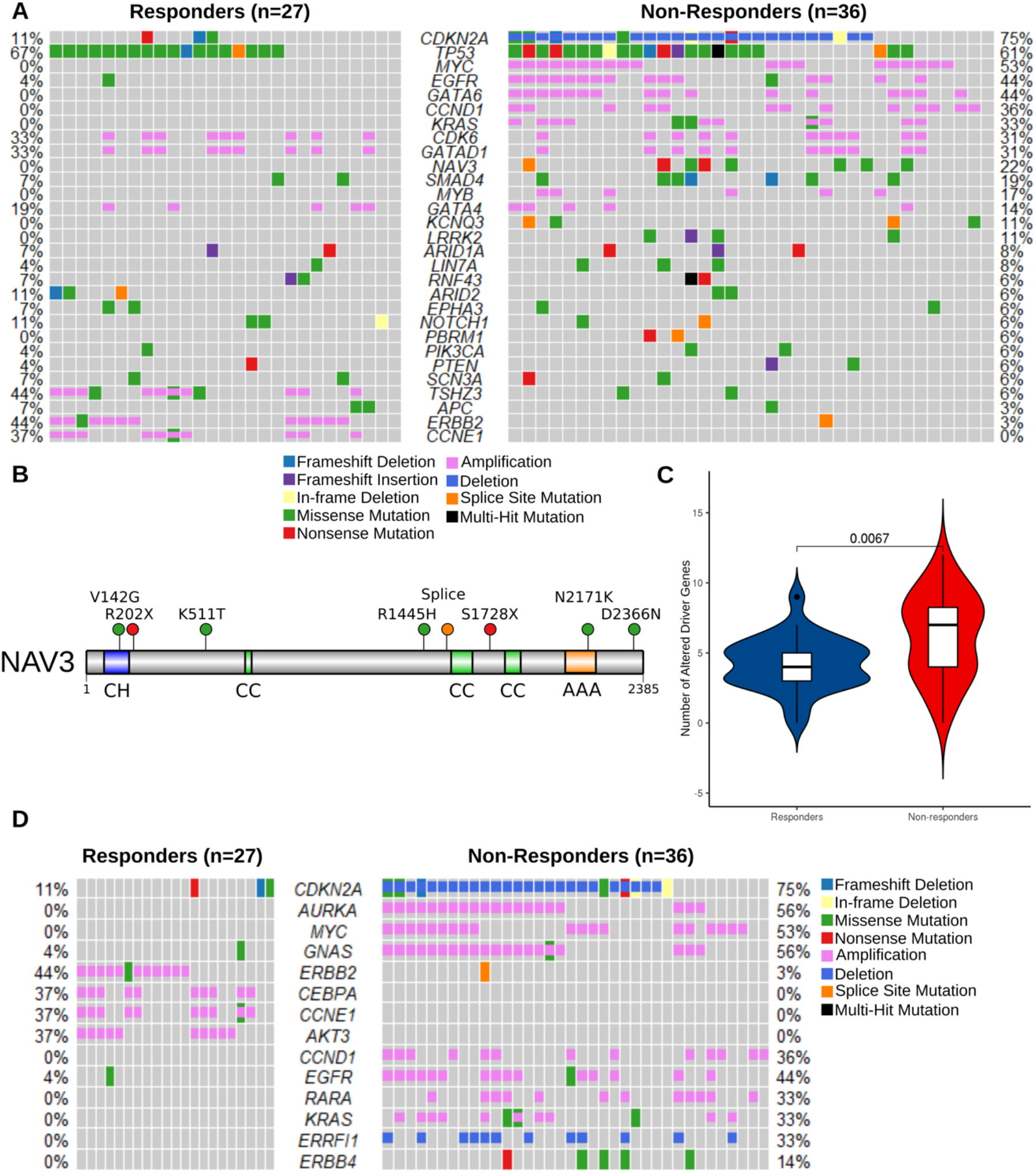
Non-responders have more EAC driver alterations of all types, including exclusive mutation of the tumor suppressor *NAV3*. (**A**) Oncoplot of SNVs, indels and CNVs combined in responders versus non-responders (n=63). Genes shown are the subset of the 76 EAC driver genes described in Frankell *et al* (2019) that were mutated in at least 5% of either group. Percentages of responders or non-responders with driver gene mutations are shown next to the corresponding row. (**B**) Protein-level diagram of mutations in the coding sequence of *NAV3*, which was exclusively mutated in nonresponders. Domains are labelled as follows: CH – Calponin Homology; CC – Coiled Coil; AAA – ATPase Associated with diverse cellular Activities. Mutational sites are shown as lollipops color-coded according to the type of mutation. (**C**) Violin plots comparing frequency of all alterations (SNVs, indels and CNVs) in EAC driver genes per sample in responders versus non-responders. (**D**) Oncoplot of SNVs, indels and CNVs combined in TARGET database genes, which are associated with a clinical action in cancer (n=63). Genes that were mutated in at least 5% of either group are shown.

We then determined which mutations might be pathogenic by cross-referencing them with the database of curated mutations (Ainscough et al. 2016). We identified 33 and 54 curated pathogenic mutations in 74% of responders and 75% of non-responders (Supplementary Tables S11-12) including three exonic mutations in *KRAS* (p.G12C, p.G12D and p.G13D). In non-responders *KRAS* was significantly co-mutated with *SMAD4* (Fisher’s exact test, two-sided, *P* = 0.006). Evidence from pancreatic cancer suggests that expression of oncogenic *KRAS* and loss of *SMAD4* cooperate to induce the expression of *EGFR* and to promote invasion (Zhao et al. 2010).

### Potentially targetable alterations in non-responders

To make sense of the complex genomic aberrations observed in this study and in EAC in general, we combined recurrent CNVs and non-synonymous mutations with the aim of identifying unique genomic aberrations in responders or non-responders. Overall, the mean number of EAC drivers carrying any alteration (SNVs, CNVs and Indels) was higher in non-responders (6.4/patient vs 4.4/patient, Wilcoxon rank sum test, *P* = 0.007, Fig. 5C). We observed that the majority of differentially altered genes were found in non-responders (Fig. 5A). Many of these genomic lesions are potentially targetable, and to investigate this further we focused on somatically altered cancer genes which are directly linked to a clinical action in the TARGET database (https://software.broadinstitute.org/cancer/cga/target) (Supplemental Table S13, S14). Non-responders displayed exclusive focal alterations of genes in this list (Fig. 5D), including: amplification of *AURKA* (16/36 non-responders, 44%); *GNAS* (20/36 nonresponders, 56%) and *RARA* (12/36 non-responders, 33%); and deletion of *ERFFI1* (12/36 nonresponders, 33%), in addition to the previously identified EAC drivers *CDKN2A, CCND1, EGFR* and *KRAS*(Supplementary Table S14). However, we observed chromosome arm amplifications of 20q, containing GNAS and AURKA, in responders. We also found potentially targetable genes exclusively amplified at the focal level in responders: *CEBPA* (13/36 responders, 37%) and *AKT3* (13/36 responders, 37%) were amplified in addition to EAC drivers *ERBB2* and *CCNE1* (Fig. 5D). These findings are in contrast to chromosome arm deletions at 19q, containing CCNE1 and CEBPA, which were found in non-responders. We examined the levels of evidence for biomarker-drug associations for our targets using the OncoKB database (Chakravarty et al. 2017). *ERBB2, EGFR* and *KRAS* have FDA-approved drugs for use in cancer therapy, whereas *CDKN2A* has biological evidence for targetability, but associated drugs are not yet standard-of-care.

## Discussion

In this study we analyzed whole-genome sequencing data from endoscopic biopsies prior to neoadjuvant chemotherapy in EAC and compared the genomes of responders to non-responders to identify potential genomic determinants of response. We comprehensively profiled CNVs, SNVs and mutational signatures in a cohort powered to identify differences between responders and non-responders. We detected distinct mutational characteristics of EAC between responders and non-responders across the spectrum from large-scale chromosomal alterations to point mutations. Our work characterises pre-existing genomic alterations that have potential as biomarkers for resistance or sensitivity to NAC.

We found that responders have higher mutational burden, in agreement with a previous published study (Findlay et al. 2016). Using a neoantigen prediction pipeline, we predicted that an increased mutational burden could lead to more abundant neoantigen recognition in responders. This could serve to bolster anti-tumor immunity as observed in the mutagenic subset of EACs reported previously (Secrier et al. 2016). Unfortunately, this study was not powered to resolve differences in NAC response between mutational signature subtypes. Although we reliably identified these mutational subtypes, there was not a clear distinction in this cohort. Consistently, we found that non-responders had impaired DNA damage response pathways and had more frequent driver gene mutations and genomic instability, despite having a lower mutation burden. The presence of an immune response related to DNA damage is known to improve survival outcomes and might contribute to this effect, as neoadjuvant therapies are genotoxic and known to stimulate anti-tumor immunity (Turkington et al. 2019).

Our data suggest that responders are dominated by point mutations, whereas non-responders display more copy number changes. Non-responders displayed a unique pattern of copy number changes characterized by chromosome arm deletions and an increased burden of copy number altered segments. This is consistent with an analysis of mutational landscapes in a pan-cancer dataset (not including EAC), which suggests that tumors are dominated by either mutations or copy number changes, but never both (Ciriello et al. 2013). The extremes of this spectrum are occupied by genomically unstable tumors, such as those observed in our cohort of non-responders. Genomic instability has been linked to a suppressed anti-tumor immunity during immunotherapy in gastric cancer (Jiang et al. 2018), whereas in non-small cell lung cancer and melanoma a higher mutation burden is linked to greater neoantigen burden (Rizvi et al. 2015) and an improved clinical response to immunotherapy (Van Allen et al. 2015; Snyder et al. 2014; Hugo et al. 2016). Taken together, this suggests that responders may be more likely to benefit from immunotherapy than non-responders and warrants further investigation.

The unique patterns of copy number changes in driver genes have important implications for treatment of chemoresistant EAC patients. These unique amplifications and deletions included potentially druggable signaling axes in EAC, and we found that MYC signaling, RTK-RAS and cell cycle pathways were preferentially mutated in non-responders. This has implications for future clinical trials, as profiling driver mutations prior to neoadjuvant treatment, such as *EGFR, CDKN2A* and *CCND1* copy number changes, would be an effective strategy to aid clinical decision-making. This would aid in the identification of patients unlikely to respond to NAC, allowing alternative or concurrent targeted therapies to be considered to exploit these pathway alterations.

Among altered pathways, we highlight cell cycle regulation as a vulnerability in non-responders. G1/S-phase checkpoint genes were disrupted, with *CCND1* and *CDK6* amplification as well as deletion of the *CDK4/6* inhibitors *CDKN2A* and *CDKN2B*. Abnormal expression of CDKs and their partner cyclins is widely reported in esophageal cancer (Kawakubo et al. 2005; Arber et al. 1999; Morgan et al. 1999), and *CCND1* amplification and nuclear expression have been shown to correlate negatively with survival (Miller et al. 2003; Bani-Hani et al. 2000). Abnormal activity of the CDK/cyclin complexes in esophageal adenocarcinoma has been shown to be a marker of acquired chemo-radio-resistance (Bani-Hani et al. 2000; Milas et al. 2002). *CDK4/6* inhibitors could be promising therapeutics for non-responders to NAC with copy number changes in this axis. In particular, *CDK4/6* inhibitors palbociclib, ribociclib and abemaciclib have shown efficacy in *in vitro* models of EAC (Frankell et al. 2019; Kosovec et al. 2017) and promising results in breast cancer, non-small cell lung cancer and melanoma patients (Patnaik et al. 2016). Similarly, the use of ABT-348, a multitarget Aurora kinase and *VEGFR* inhibitor (Maitland et al. 2018), is currently being explored in phase I and II clinical trials in patients with *CDKN2A*-deficient tumors (Sharma 2015; Hong 2015), suggesting additional targeted therapies to this axis are closer to clinical adoption.

Previous genomic analyses suggest that copy number changes to RTKs are pervasive in EAC, with the potential for targeting with RTK inhibitors specific to the activated pathways, such as trastuzumab and ABT-806 for *ERBB2* and *EGFR* respectively (Secrier et al. 2016; Catenacci et al. 2020). We found that *EGFR* is uniquely amplified in non-responders, suggesting that anti-*EGFR* antibodies cetuximab or ABT-806 may be useful therapies in these patients. Cetuximab is well tolerated by EAC patients (Chan et al. 2011) and has gone through phase III trials as a neoadjuvant therapy in addition to chemotherapy or chemoradiation, showing a modest improvement in recurrence-free survival in an unselected population (Ruhstaller et al. 2018). *EGFR*-amplified EAC patients have been shown to particularly benefit from cetuximab (Luber et al. 2011). However, *KRAS* mutations are known to confer resistance to cetuximab in colorectal cancer (Van Cutsem et al. 2009; Lièvre et al. 2006), and while this is unclear in EAC due to the rarity of *KRAS* mutations and unselected patient populations (Ruhstaller et al. 2018; Richards et al. 2013), *KRAS*-mutant tumors in our dataset bore the same mutations in codons 12 and 13 as the resistant colorectal tumors and were non-responders. This underscores the need for careful selection of patient populations to be treated with RTK inhibitors based on *KRAS* mutation status.

Finally, we report that Neuron Navigator-3 (*NAV3*), a known tumor suppressor downstream of *EGFR* (Cohen-Dvashi et al. 2015), is mutated exclusively in non-responders. *NAV3* is a microtubule-binding protein whose expression is regulated by *TP73* and induced by *EGF* in breast cancer cells (Cohen-Dvashi et al. 2015). In our cohort we observed that *NAV3* was co-mutated with *CDKN2A*, with most mutations being missense. Predictions of neoantigen recognition in lung adenocarcinoma suggest that *NAV3* is one of the most commonly mutated genes with predicted neoantigen recognition in this disease as well (Cai et al. 2018). The functional consequences of these mutations in EAC are unclear but we predict several to be functionally deleterious.

*NAV3* is primarily implicated as a metastasis suppressor in multiple cancer types*. NAV3* is upregulated in response to DNA damage in colon carcinoma cells and is involved in the suppression of migration and invasion *in vitro* (Uboveja et al. 2020). Loss of heterozygosity occurs in colorectal cancer and this associates with lymph node metastasis (Carlsson et al. 2012). *NAV3* expression is attenuated in metastatic colon cancer (Uboveja et al. 2020), breast cancer and lung cancer (Cohen-Dvashi et al. 2015) and its knockdown promotes invasive behaviours (Cohen-Dvashi et al. 2015; Uboveja et al. 2020), platinum drug resistance (Pink et al. 2015) and epithelial mesenchymal transition *in vitro* (Uboveja et al. 2020) and enhances metastasis *in vivo* (Cohen-Dvashi et al. 2015). The inhibitory effect of *NAV3* on invasion and metastasis may be due to its promotion of slower, directional cell migration as opposed to the random migration observed in *NAV3* knockdown cells, which enhances their ability to explore their environment (Cohen-Dvashi et al. 2015). Silencing of *NAV3 in vitro* also leads to upregulation of *IL-23R* in colorectal (Carlsson et al. 2012) and glioma cell lines (Carlsson et al. 2013), linked to proinflammatory JAK-STAT signalling. A sizeable proportion of EAC non-responders (22%) carry mutations in *NAV3*, and its status as a unique genetic lesion to this group suggests that *NAV3* mutation could be used as a biomarker to identify some of the patients who fail to respond to NAC. This warrants validation in a larger cohort of patients, including further study of *NAV3* expression and the functional consequences of *NAV3* mutation in EAC.

This study is not without shortcomings. With 63 patients we were able to resolve genomic differences between responders and non-responders at the copy number and mutational level, but had insufficient sample size to fully study the impact of mutational signatures on NAC response. As EACs accrue many genetic alterations and very few are recurrent (Frankell et al. 2019), we lack the power to resolve the significance of rarer mutations on survival and to determine rarer co-mutations. However, even with limited sample size, WGS was able to identify KRAS mutations in non-responders, which are frequently associated with treatment resistance in colorectal cancer (Van Cutsem et al. 2009; Lièvre et al. 2006), demonstrating the robustness of our approach. Despite these shortcomings, our dataset of responders and non-responders to NAC is the largest of its kind in EAC and represents a step forward in our understanding of the genetic determinants of NAC resistance.

In summary, we identified genetic features and mutations that are uniquely associated with response to NAC. This indicates the presence of a subset of patients harboring pre-existing mutations that confer resistance to NAC. Importantly, these mutations are potentially clinically actionable, with a variety of drugs in clinical trials to support a targeted therapy strategy - an approach that has previously met with success in metastatic EAC patients (Catenacci et al. 2020). We envision a treatment pipeline that incorporates driver mutation profiling in EAC, combining response prediction with targeted therapies to enhance response to NAC and improve survival outcomes.

## Methods

### Overview of patients and sequencing strategy

EAC patients in this study are presented in Figure 1A & Table 1. Sample collection and processing were performed as previously described (Noorani et al. 2017) as part of the OCCAMS (Oesophageal Cancer Clinical and Molecular Stratification) Consortium. Pathological tumor response was assessed in the resection specimens by tumor regression grading (TRG) (Mandard et al. 1994) with responders defined as TRG ≤2 and non-responders as TRG ≥4. Mandard grade was scored by a specialist gastrointestinal pathologist blinded to the clinical data at the treating cancer centre.

### Whole-genome sequencing analysis

WGS, single nucleotide variant (SNV) and small insertion or deletion (Indel) calling has been performed using Strelka (Saunders et al. 2012) (version 2.0.15) against the GRCh37 reference genome as described by Secrier et al. (2016), with 94% of the known genome being sequenced while achieving a PHRED quality of at least 30 for at least 80% of mapping bases.

Functional annotation of the resulting variants was performed using ANNOVAR (Yang and Wang 2015) and the Ensembl Variant Effect Predictor (VEP) (https://www.ensembl.org/Tools/VEP). Furthermore, 536 false positive genes (Mourikis et al. 2019) were removed from subsequent analysis. Data visualisation including oncoprints and lollipop plots were performed by maftools (version 2.4.15) (Mayakonda et al. 2018). Mutually exclusiveness or co-occurrence analysis of genes was also performed by maftools using pair-wise Fisher’s Exact test to detect significant pairs of given genes.

### Copy number and clonality analysis

Absolute genome copy number following correction for estimated normal-cell contamination was called using ASCAT package (version 2.3) in R (Van Loo et al. 2010). Cellularity and ploidy estimates were also obtained using ASCAT and samples with estimated cellularity <20% were removed from further analysis. Significantly amplified/deleted regions in the cohort were identified using GISTIC2.0 (Mermel et al. 2011). Copy number variations (CNVs) were corrected for ploidy (= total copy number of the segment / average estimated ploidy of each sample) and GISTIC was run on an input defined as the log2 of the CNV with gain (-ta) >= 1.0 and loss (-td) <= 0.4 respectively.

### Genomic instability analysis

Copy number burden represents the fraction of bases deviating from baseline ploidy (defined as above 0.5 or below - 0.5 in log2 relative copy number space and in segments > 1kb length) named as genomic instability score (GII). CIN70 score was calculated by averaging the FPKM expression values of CIN70 signature in each sample from available RNA-seq data.

### Mutational signature and neoantigen analysis

Tumor Mutation Burden (TMB) in terms of per megabases was measured with 50 MB capture size for non-synonymous mutations.

In order to compare the neoantigen load between two groups, we used binding affinity for patient-specific class I human lymphocyte antigen (HLA) alleles, constituting potential candidate neoantigens by checking for the binding strength for peptides of length 9 using NeoPredPipe (Schenck et al. 2019). We then quantified the peptides that displayed high affinity (recognition potential > 1) binding in tumor, but no binding in the respective matched normal (Luksza et al. 2017) as recognition potential prediction step implemented in NeoPredPipe to obtain recognition potential for each sample. We then compared the recognition potential between two groups by Wilcoxon rank sum test. EAC mutational signatures were extracted using SigProfiler. We have processed the COSMIC solution to remove any artefactual signatures and signatures that contribute on average less than 1% of the mutations in the genome. A total of nine mutational signatures were identified, of which six have been previously identified in EAC described by Secrier et al. (2016): SBS17A and SBS17B dominated by T>G substitutions in a CTT context and possibly associated with gastric acid reflux; SBS3, a complex pattern caused by defects in the BRCA1/2-led homologous recombination pathway; SBS2, C>T mutations in a TCA/TCT context, an APOBEC-driven hypermutated phenotype; SBS1, C>T in a *CG context, associated with aging processes; An SBS18-like signature, C>A/T dominant in a GCA/TCT context, formerly described in neuroblastoma, breast and stomach cancers; SBS13 is usually found in the same samples as SBS2; SBS5, linked to tobacco exposure; and SBS41 with unknown aetiology. Clustering of mutational signatures was performed with the NMF package (version 0.23.0) (Gaujoux and Seoighe 2010) set to three main clusters as previously observed in EAC.

### Expression profiling by bulk RNA sequencing (RNA-seq) and Gene Set Enrichment Analysis (GSEA)

We were able to explore gene expression changes to investigate the expression levels of recurrently amplified/deleted genes, between responders and non-responders in 30 RNA-seq samples matched with WGS data (responders/9, non-responders/21). For a given EAC known driver identified as recurrently amplified or deleted in either group, we compared Fragments Per Kilobase of transcript, per Million mapped reads (FPKM) values for that gene by Wilcoxon rank sum test. For GSEA by using MSigDB hallmark gene sets, we used normalized values from DESeq2 as input. We used GSEA and DESeq2 modules both implemented online (https://www.gsea-msigdb.org/gsea/) (Subramanian et al. 2005) with default parameters.

### Survival analysis

For relating CNVs to overall survival, we used a Boolean matrix of CNV status of EAC driver genes. Multivariate analyses were performed by Cox proportional hazards regression model using the survival package (version 3.2.7) in R. Overall survival was defined as the time interval from initial surgical excision to death or last follow-up time (censored) and Kaplan-Meier plots were visualised using the ggkm 1.0 R package (https://github.com/michaelway/ggkm).

### DDR pathway deregulation analysis

The Pathifier algorithm (version v1.0) calculates for any given pathway a deregulation score (PDS) for each cancer sample, based on gene expression data (log2 normalized) (Drier et al. 2013). Only the 5000 genes with the largest variation over available RNA-seq samples were used as input to the algorithm. PDS score represents the extent to which the activity of the pathway differs in a particular sample from the activity in the opponent sample. Here responders and non-responders were used as opponent groups of samples. We calculated an average of PDS over sixteen DDR sub-pathways by using more than 450 genes associated with DDR, as previously described in a pan-cancer analysis (Pearl et al. 2015).

### Classification of genes relevant for genomics-driven therapy

To identify genes relevant for genomics-driven therapy, we used version 2.0 of TARGET (tumor alterations relevant for genomics-driver therapy) database (www.broadinstitute.org/cancer/cga/target). We also used OncoKB (Chakravarty et al. 2017)(https://www.oncokb.org/) for the association of drug-biomarkers of differentially mutated genes.

### Identification of specific mutations with therapeutic relevance

The DoCM (Ainscough et al. 2016) was used to identify mutations with clinical evidence (drug targets associated with a mutation; diagnostic or prognostic markers associated with a mutation) or functional evidence (disease function described in cell lines; disease function described in animal models). The database is available online at docm.genome.wustl.edu.

### Statistics

Measurements between groups were compared using the Wilcoxon rank-sum test for continuous data with non-normal distribution and T-test for data with normal distribution or Fisher’s exact test for count data.

### Data Access

The WGS and RNA-seq data generated in this study are available from the European Genome-Phenome Archive (https://ega-archive.org/) under accession number EGAD00001007493.

### Members of Oesophageal Cancer Clinical and Molecular Stratification (OCCAMS) Consortium

Rebecca C. Fitzgerald^1^, Paul A.W. Edwards^1,2^, Nicola Grehan^1^, Barbara Nutzinger^1^, Elwira Fidziukiewicz^1^, Shona MacRae^1^, Aisling M Redmond^1^, Alex Northrop^1^, Gianmarco Contino^1^, Annalise Katz-Summercorn^1^, Sujath Abbas^1^, Elizabeth C. Smyth^5^, Maria O’Donovan^1,3^, Ahmad Miremadi^1,3^, Shalini Malhotra^1,3^, Monika Tripathi^1,3^, Matthew Eldridge^2^, Maria Secrier^2^, Ginny Devonshire^2^, Sriganesh Jammula^2^, Jim Davies^4^, Charles Crichton^4^, Nick Carroll^5^, Peter Safranek^5^, Andrew Hindmarsh^5^, Vijayendran Sujendran^5^, Stephen J. Hayes^6,13^, Yeng Ang^6,7,26^, Andrew Sharrocks^26^, Shaun R. Preston^8^, Izhar Bagwan^8^, Vicki Save^9^, Richard J.E. Skipworth^9^, Ted R. Hupp^20^, J. Robert O’Neill^5^,^9^,^20^, Olga Tucker^10,29^, Andrew Beggs^10,25^, Philippe Taniere^10^, Sonia Puig^10^, Timothy J. Underwood^11,12^, Robert C. Walker^11,12^, Ben L. Grace^11^, Jesper Lagergren^14,22^, James Gossage^14,21^, Andrew Davies^14,21^, Fuju Chang^14,21^, Janine Zylstra^14,21^, Ula Mahadeva^14^, Vicky Goh^21^, Francesca D. Ciccarelli^21^, Grant Sanders^15^, Richard Berrisford^15^, David Chan^15^, Mike Lewis^16^, Ed Cheong^16^, Bhaskar Kumar^16^, L. Sreedharan^16^ Simon L Parsons^17^, Irshad Soomro^17^, Philip Kaye^17^, John Saunders^6^, Laurence Lovat^18^, Rehan Haidry^18^, Michael Scott^19^, Sharmila Sothi^23^, Suzy Lishman^2^, George B. Hanna^27^, Christopher J. Peters^27^,Krishna Moorthy^27^, Anna Grabowska^28^, Richard Turkington^30^, Damian McManus^30^, Helen Coleman^30^, David Khoo^31^, Will Fickling^31^, Russell D Petty^32^

^1^ Medical Research Council Cancer Unit, Hutchison/Medical Research Council Research Centre, University of Cambridge, Cambridge, UK

^2^ Cancer Research UK Cambridge Institute, University of Cambridge, Cambridge, UK

^3^ Department of Histopathology, Addenbrooke’s Hospital, Cambridge, UK

^4^ Department of Computer Science, University of Oxford, UK, OX1 3QD

^5^ Cambridge University Hospitals NHS Foundation Trust, Cambridge, UK, CB2 0QQ

^6^ Salford Royal NHS Foundation Trust, Salford, UK, M6 8HD

^7^ Wigan and Leigh NHS Foundation Trust, Wigan, Manchester, UK, WN1 2NN

^8^ Royal Surrey County Hospital NHS Foundation Trust, Guildford, UK, GU2 7XX

^9^ Edinburgh Royal Infirmary, Edinburgh, UK, EH16 4SA

^10^ University Hospitals Birmingham NHS Foundation Trust, Birmingham, UK, B15 2GW

^11^ University Hospital Southampton NHS Foundation Trust, Southampton, UK, SO16 6YD

^12^ Cancer Sciences Division, University of Southampton, Southampton, UK, SO17 1BJ

^13^ Faculty of Medical and Human Sciences, University of Manchester, UK, M13 9PL

^14^ Guy’s and St Thomas’s NHS Foundation Trust, London, UK, SE1 7EH

^15^ Plymouth Hospitals NHS Trust, Plymouth, UK, PL6 8DH

^16^ Norfolk and Norwich University Hospital NHS Foundation Trust, Norwich, UK, NR4 7UY

^17^ Nottingham University Hospitals NHS Trust, Nottingham, UK, NG7 2UH

^18^ University College London, London, UK, WC1E 6BT

^19^ Wythenshawe Hospital, Manchester, UK, M23 9LT

^20^ Edinburgh University, Edinburgh, UK, EH8 9YL

^21^ King’s College London, London, UK, WC2R 2LS

^22^ Karolinska Institute, Stockholm, Sweden, SE-171 77

^23^ University Hospitals Coventry and Warwickshire NHS, Trust, Coventry, UK, CV2 2DX

^24^ Peterborough Hospitals NHS Trust, Peterborough City Hospital, Peterborough, UK, PE3 9GZ

^25^ Institute of Cancer and Genomic sciences, University of Birmingham, B15 2TT

^26^ GI science centre, University of Manchester, UK, M13 9PL.

^27^ Department of Surgery and Cancer, Imperial College, London, UK, W2 1NY

^28^ Queen’s Medical Centre, University of Nottingham, Nottingham, UK

^29^ Heart of England NHS Foundation Trust, Birmingham, UK, B9 5SS.

^30^ Centre for Cancer Research and Cell Biology, Queen’s University Belfast, Northern Ireland, UK, BT7 1NN.

^31^ Queen’s Hospital, Romford, RM7 0AG

^32^ Tayside Cancer Centre, Ninewells Hospital and Medical School, Dundee, DD1 9SY

## Acknowledgements

This work was supported by Cancer Research UK & Royal College of Surgeons of England Advanced Clinician Scientist Fellowship to T.J.U. “Cellular interplay in oesophageal cancer.” A23924. M.J.R.Z was supported by the Wessex Medical Research Innovation Fund 2011 R06 and 2014 U10, Leuka Charity, John Goldman Fellowship 2016/JGF/0003 and the Southampton CRUK Centre Development Fund. Sample sequencing was funded through the Oesophageal Cancer Clinical and Molecular Stratification (OCCAMS) Consortium as part of the International Cancer Genome Consortium, and was funded by a programme grant from Cancer Research UK (RG66287). OCCAMS2 was funded by a Programme Grant from Cancer Research UK (RG81771/84119). The laboratory of R.C.F. is funded by a Core Programme Grant from the Medical Research Council (RG84369). This research was supported by the NIHR Cambridge Biomedical Research Centre (BRC-1215-20014). The views expressed are those of the authors and not necessarily those of the NIHR or the Department of Health and Social Care.

## Author Contributions

Author Contributions: Conceptualization, F.I., M.J.J.R-Z., T.J.U; Methodology, F.I., G.D., M.S. Formal analysis, F.I., M.S.; Investigation, F.I.; Data Curation, F.I., G.D.; Writing – Original Draft, F.I., B.S., S.B.; Writing – Review & Editing, M.J.J.R-Z., B.S., S.B., M.S.; Visualization, F.I, B.S., S.B.; Supervision, M.J.J.R-Z., T.J.U., R.C.F.; Project Administration, F.I., M.J.J.R-Z., T.J.U; Funding Acquisition, T.J.U, R.C.F.;

## References

Adzhubei IA, et al. A method and server for predicting damaging missense mutations. Nat Methods. 2010;7(4):248–9.

Ainscough BJ, Griffith M, Coffman AC, Wagner AH, Kunisaki J, Choudhary MNK, McMichael JF, Fulton RS, Wilson RK, Griffith OL, et al. 2016. DoCM: A database of curated mutations in cancer. Nat Methods.

Al-Batran SE, Homann N, Pauligk C, Goetze TO, Meiler J, Kasper S, Kopp HG, Mayer F, Haag GM, Luley K, et al. 2019. Perioperative chemotherapy with fluorouracil plus leucovorin, oxaliplatin, and docetaxel versus fluorouracil or capecitabine plus cisplatin and epirubicin for locally advanced, resectable gastric or gastro-oesophageal junction adenocarcinoma (FLOT4): a randomised, phase 2/3 trial. Lancet 393: 1948–1957. https://pubmed.ncbi.nlm.nih.gov/30982686/ (Accessed January 14, 2021).

Alexandrov LB, Kim J, Haradhvala NJ, Huang MN, Tian Ng AW, Wu Y, Boot A, Covington KR, Gordenin DA, Bergstrom EN, et al. 2020. The repertoire of mutational signatures in human cancer. Nature.

Allum WH, Stenning SP, Bancewicz J, Clark PI, Langley RE. 2009. Long-term results of a randomized trial of surgery with or without preoperative chemotherapy in esophageal cancer. J Clin Oncol 27: 5062–5067. https://pubmed.ncbi.nlm.nih.gov/19770374/ (Accessed January 14, 2021).

Arber N, Gammon MD, Hibshoosh H, Britton JA, Zhang Y, Schonberg JB, Roterdam H, Fabian I, Holt PR, Weinstein IB. 1999. Overexpression of cyclin D1 occurs in both squamous carcinomas and adenocarcinomas of the esophagus and in adenocarcinomas of the stomach. Hum Pathol.

Bani-Hani K, Martin IG, Hardie LJ, Mapstone N, Briggs JA, Forman D, Wild CP. 2000. Prospective study of cyclin D1 overexpression in Barrett’s esophagus: Association with increased risk of adenocarcinoma. J Natl Cancer Inst.

Cai W, Zhou D, Wu W, Tan WL, Wang J, Zhou C, Lou Y. 2018. MHC class II restricted neoantigen peptides predicted by clonal mutation analysis in lung adenocarcinoma patients: Implications on prognostic immunological biomarker and vaccine design. BMC Genomics 19. https://pubmed.ncbi.nlm.nih.gov/30075702/ (Accessed January 19, 2021).

Carlsson E, Krohn K, Ovaska K, Lindberg P, Häyry V, Maliniemi P, Lintulahti A, Korja M, Kivisaari R, Hussein S, et al. 2013. Neuron navigator 3 alterations in nervous system tumors associate with tumor malignancy grade and prognosis. Genes Chromosom Cancer 52: 191–201. https://pubmed.ncbi.nlm.nih.gov/23097141/ (Accessed January 19, 2021).

Carlsson E, Ranki A, Sipilä L, Karenko L, Abdel-Rahman WM, Ovaska K, Siggberg L, Aapola U, Ssämäki R, Häyry V, et al. 2012. Potential role of a navigator gene NAV3 in colorectal cancer. Br J Cancer 106: 517–524. https://pubmed.ncbi.nlm.nih.gov/22173670/ (Accessed January 19, 2021).

Carter SL, Eklund AC, Kohane IS, Harris LN, Szallasi Z. 2006. A signature of chromosomal instability inferred from gene expression profiles predicts clinical outcome in multiple human cancers. Nat Genet.

Catenacci DVT, Moya S, Lomnicki S, Chase LM, Peterson BF, Reizine N, Alpert L, Setia N, Xiao S-Y, Hart J, et al. 2020. Personalized antibodies for gastroesophageal adenocarcinoma (PANGEA): a phase 2 study evaluating an individualized treatment strategy for metastatic disease. Cancer Discov CD-20-1408. http://cancerdiscovery.aacrjournals.org/lookup/doi/10.1158/2159-8290.CD-20-1408 (Accessed November 25, 2020).

Chakiba C, Lagarde P, Pissaloux D, Neuville A, Brulard C, Pérot G, Coindre JM, Terrier P, Ranchere-Vince D, Ferrari A, et al. 2014. Response to chemotherapy is not related to chromosome instability in synovial sarcoma. Ann Oncol.

Chakravarty D, Gao J, Phillips S, Kundra R, Zhang H, Wang J, Rudolph JE, Yaeger R, Soumerai T, Nissan MH, et al. 2017. OncoKB: A Precision Oncology Knowledge Base. JCO Precis Oncol 2017: 1–16. https://pubmed.ncbi.nlm.nih.gov/28890946/ (Accessed January 14, 2021).

Chan JA, Blaszkowsky LS, Enzinger PC, Ryan DP, Abrams TA, Zhu AX, Temel JS, Schrag D, Bhargava P, Meyerhardt JA, et al. 2011. A multicenter phase II trial of single-agent cetuximab in advanced esophageal and gastric adenocarcinoma. Ann Oncol 22: 1367–1373. https://pubmed.ncbi.nlm.nih.gov/21217058/ (Accessed January 14, 2021).

Ciriello G, Miller ML, Aksoy BA, Senbabaoglu Y, Schultz N, Sander C. 2013. Emerging landscape of oncogenic signatures across human cancers. Nat Genet.

Cohen-Dvashi H, Ben-Chetrit N, Russell R, Carvalho S, Lauriola M, Nisani S, Mancini M, Nataraj N, Kedmi M, Roth L, et al. 2015. Navigator-3, a modulator of cell migration, may act as a suppressor of breast cancer progression. EMBO Mol Med 7: 299–314.

Cramer G, Simone CB, Busch TM, Cengel KA. 2018. Adjuvant, neoadjuvant, and definitive radiation therapy for malignant pleural mesothelioma. J Thorac Dis.

Cunningham D, Allum WH, Stenning SP, Thompson JN, Van de Velde CJH, Nicolson M, Scarffe JH, Lofts FJ, Falk SJ, Iveson TJ, et al. 2006. Perioperative Chemotherapy versus Surgery Alone for Resectable Gastroesophageal Cancer. N Engl J Med 355: 11–20. https://pubmed.ncbi.nlm.nih.gov/16822992/ (Accessed January 14, 2021).

Do Canto LM, Larsen SJ, Kupper BEC, De Souza Begnami MDF, Scapulatempo-Neto C, Petersen AH, Aagaard MM, Baumbach J, Aguiar S, Rogatto SR. 2019. Increased levels of genomic instability and mutations in homologous recombination genes in locally advanced rectal carcinomas. Front Oncol 9. https://pubmed.ncbi.nlm.nih.gov/31192117/ (Accessed January 15, 2021).

Drier Y, Sheffer M, Domany E. 2013. Pathway-based personalized analysis of cancer. Proc Natl Acad Sci U S A 110: 6388–6393. www.pnas.org/cgi/doi/10.1073/pnas.1219651110 (Accessed February 12, 2021).

Findlay JM, Castro-Giner F, Makino S, Rayner E, Kartsonaki C, Cross W, Kovac M, Ulahannan D, Palles C, Gillies RS, et al. 2016. Differential clonal evolution in oesophageal cancers in response to neo-adjuvant chemotherapy. Nat Commun 7.

Frankell AM, Jammula SG, Li X, Contino G, Killcoyne S, Abbas S, Perner J, Bower L, Devonshire G, Ococks E, et al. 2019. The landscape of selection in 551 esophageal adenocarcinomas defines genomic biomarkers for the clinic. Nat Genet 51: 506–516.

Gaujoux R, Seoighe C. 2010. A flexible R package for nonnegative matrix factorization. BMC Bioinformatics.

Girling DJ, Bancewicz J, Clark PI, Smith DB, Donnelly RJ, Fayers PM, Weeden S, Girling DJ, Hutchinson T, Harvey A, et al. 2002. Surgical resection with or without preoperative chemotherapy in oesophageal cancer: A randomised controlled trial. Lancet 359: 1727–1733. https://pubmed.ncbi.nlm.nih.gov/12049861/ (Accessed January 14, 2021).

Greenbaum A, Martin DR, Bocklage T, Lee JH, Ness SA, Rajput A. 2019. Tumor Heterogeneity as a Predictor of Response to Neoadjuvant Chemotherapy in Locally Advanced Rectal Cancer. Clin Colorectal Cancer 18: 102–109.

Höglander EK, Nord S, Wedge DC, Lingjærde OC, Silwal-Pandit L, Gythfeldt HVL, Vollan HKM, Fleischer T, Krohn M, Schlitchting E, et al. 2018. Time series analysis of neoadjuvant chemotherapy and bevacizumab-treated breast carcinomas reveals a systemic shift in genomic aberrations. Genome Med.

Hong DS. 2015. Phase II Study of Ilorasertib (ABT348) in Patients With CDKN2A Deficient Solid Tumors - Full Text View - ClinicalTrials.gov. https://clinicaltrials.gov/ct2/show/NCT02478320 (Accessed January 19, 2021).

Hugo W, Zaretsky JM, Sun L, Song C, Moreno BH, Hu-Lieskovan S, Berent-Maoz B, Pang J, Chmielowski B, Cherry G, et al. 2016. Genomic and Transcriptomic Features of Response to Anti-PD-1 Therapy in Metastatic Melanoma. Cell.

Jiang Z, Liu Z, Li M, Chen C, Wang X. 2018. Immunogenomics Analysis Reveals that TP53 Mutations Inhibit Tumor Immunity in Gastric Cancer. Transl Oncol.

Kawakubo H, Ozawa S, Ando N, Kitagawa Y, Mukai M, Ueda M, Kitajima M. 2005. Alterations of p53, cyclin D1 and pRB expression in the carcinogenesis of esophageal squamous cell carcinoma. Oncol Rep.

Kosovec JE, Zaidi AH, Omstead AN, Matsui D, Biedka MJ, Cox EJ, Campbell PT, Biederman RWW, Kelly RJ, Jobe BA. 2017. CDK4/6 dual inhibitor abemaciclib demonstrates compelling preclinical activity against esophageal adenocarcinoma: A novel therapeutic option for a deadly disease. Oncotarget 8: 100421–100432. https://pubmed.ncbi.nlm.nih.gov/29245989/ (Accessed January 14, 2021).

Lesurf R, Griffith OL, Griffith M, Hundal J, Trani L, Watson MA, Aft R, Ellis MJ, Ota D, Suman VJ, et al. 2017. Genomic characterization of HER2-positive breast cancer and response to neoadjuvant trastuzumab and chemotherapy-results from the ACOSOG Z1041 (Alliance) trial. Ann Oncol Off J Eur Soc Med Oncol.

Li X, Francies HE, Secrier M, Perner J, Miremadi A, Galeano-Dalmau N, Barendt WJ, Letchford L, Leyden GM, Goffin EK, et al. 2018. Organoid cultures recapitulate esophageal adenocarcinoma heterogeneity providing a model for clonality studies and precision therapeutics. Nat Commun 9.

Li Z, Gao X, Peng X, Chen MJM, Li Z, Wei B, Wen X, Wei B, Dong Y, Bu Z, et al. 2020. Multi-omics characterization of molecular features of gastric cancer correlated with response to neoadjuvant chemotherapy. Sci Adv.

Lièvre A, Bachet JB, Le Corre D, Boige V, Landi B, Emile JF, Côté JF, Tomasic G, Penna C, Ducreux M, et al. 2006. KRAS mutation status is predictive of response to cetuximab therapy in colorectal cancer. Cancer Res 66: 3992–3995. https://pubmed.ncbi.nlm.nih.gov/16618717/ (Accessed January 14, 2021).

Luber B, Deplazes J, Keller G, Walch A, Rauser S, Eichmann M, Langer R, Höfler H, Hegewisch-Becker S, Folprecht G, et al. 2011. Biomarker analysis of cetuximab plus oxaliplatin/leucovorin/5-fluorouracil in first-line metastatic gastric and oesophago-gastric junction cancer: results from a phase II trial of the Arbeitsgemeinschaft Internistische Onkologie (AIO). BMC Cancer 11. https://pubmed.ncbi.nlm.nih.gov/22152101/ (Accessed January 14, 2021).

Luksza M, Riaz N, Makarov V, Balachandran VP, Hellmann MD, Solovyov A, Rizvi NA, Merghoub T, Levine AJ, Chan TA, et al. 2017. A neoantigen fitness model predicts tumour response to checkpoint blockade immunotherapy. Nature.

Maitland ML, Piha-Paul S, Falchook G, Kurzrock R, Nguyen L, Janisch L, Karovic S, McKee M, Hoening E, Wong S, et al. 2018. Clinical pharmacodynamic/exposure characterisation of the multikinase inhibitor ilorasertib (ABT-348) in a phase 1 dose-escalation trial. Br J Cancer 118: 1042–1050. https://doi.org/10.1038/s41416-018-0020-2 (Accessed January 15, 2021).

Mancini R, Pattaro G, Diodoro MG, Sperduti I, Garufi C, Stigliano V, Perri P, Grazi GL, Cosimelli M. 2018. Tumor Regression Grade After Neoadjuvant Chemoradiation and Surgery for Low Rectal Cancer Evaluated by Multiple Correspondence Analysis: Ten Years as Minimum Follow-up. Clin Colorectal Cancer.

Mandard A-M, Dalibard F, Mandard J-C, Marnay J, Henry-Amar M, Petiot J-F, Roussel A, Jacob J-H, Segol P, Samama G, et al. 1994. Pathologic assessment of tumor regression after preoperative chemoradiotherapy of esophageal carcinoma. Clinicopathologic correlations. Cancer.

Mayakonda A, Lin DC, Assenov Y, Plass C, Koeffler HP. 2018. Maftools: Efficient and comprehensive analysis of somatic variants in cancer. Genome Res.

McLaren W, Gil L, Hunt SE, Riat HS, Ritchie GRS, Thormann A, Flicek P, Cunningham F. 2016. The Ensembl Variant Effect Predictor. Genome Biol 17: 122. http://genomebiology.biomedcentral.com/articles/10.1186/s13059-016-0974-4 (Accessed January 14, 2021).

Mermel CH, Schumacher SE, Hill B, Meyerson ML, Beroukhim R, Getz G. 2011. GISTIC2.0 facilitates sensitive and confident localization of the targets of focal somatic copy-number alteration in human cancers. Genome Biol 12. https://pubmed.ncbi.nlm.nih.gov/21527027/ (Accessed January 14, 2021).

Milas L, Akimoto T, Hunter NR, Mason KA, Buchmiller L, Yamakawa M, Muramatsu H, Ang KK. 2002. Relationship between cyclin D1 expression and poor radioresponse of murine carcinomas. Int J Radiat Oncol Biol Phys.

Miller CT, Moy JR, Lin L, Schipper M, Normolle D, Brenner DE, Iannettoni MD, Orringer MB, Beer DG. 2003. Gene Amplification in Esophageal Adenocarcinomas and Barrett’s with High-Grade Dysplasia. Clin Cancer Res.

Morgan RJ, Newcomb P V., Hardwick RH, Alderson D. 1999. Amplification of cyclin D1 and MDM-2 in oesophageal carcinoma. Eur J Surg Oncol.

Mourikis TP, Benedetti L, Foxall E, Temelkovski D, Nulsen J, Perner J, Cereda M, Lagergren J, Howell M, Yau C, et al. 2019. Patient-specific cancer genes contribute to recurrently perturbed pathways and establish therapeutic vulnerabilities in esophageal adenocarcinoma. Nat Commun.

Murugaesu N, Wilson GA, Birkbak NJ, Watkins TBK, McGranahan N, Kumar S, Abbassi-Ghadi N, Salm M, Mitter R, Horswell S, et al. 2015. Tracking the genomic evolution of esophageal adenocarcinoma through neoadjuvant chemotherapy. Cancer Discov 5: 821–832.

Noble F, Lloyd MA, Turkington R, Griffiths E, O’Donovan M, O’Neill JR, Mercer S, Parsons SL, Fitzgerald RC, Underwood TJ, et al. 2017. Multicentre cohort study to define and validate pathological assessment of response to neoadjuvant therapy in oesophagogastric adenocarcinoma. Br J Surg.

Noble F, Mellows T, McCormick Matthews LH, Bateman AC, Harris S, Underwood TJ, Byrne JP, Bailey IS, Sharland DM, Kelly JJ, et al. 2016. Tumour infiltrating lymphocytes correlate with improved survival in patients with oesophageal adenocarcinoma. Cancer Immunol Immunother 65: 651–662. https://www.ncbi.nlm.nih.gov/pubmed/27020682.

Nones K, Waddell N, Wayte N, Patch AM, Bailey P, Newell F, Holmes O, Fink JL, Quinn MCJ, Tang YH, et al. 2014. Genomic catastrophes frequently arise in esophageal adenocarcinoma and drive tumorigenesis. Nat Commun.

Noorani A, Bornschein J, Lynch AG, Secrier M, Achilleos A, Eldridge M, Bower L, Weaver JMJ, Crawte J, Ong CA, et al. 2017. A comparative analysis of whole genome sequencing of esophageal adenocarcinoma pre-and post-chemotherapy. Genome Res 27: 902–912.

Patnaik A, Rosen LS, Tolaney SM, Tolcher AW, Goldman JW, Gandhi L, Papadopoulos KP, Beeram M, Rasco DW, Hilton JF, et al. 2016. Efficacy and safety of Abemaciclib, an inhibitor of CDK4 and CDK6, for patients with breast cancer, non–small cell lung cancer, and other solid tumors. Cancer Discov 6: 740–753. https://pubmed.ncbi.nlm.nih.gov/27217383/ (Accessed January 14, 2021).

Pearl LH, Schierz AC, Ward SE, Al-Lazikani B, Pearl FMG. 2015. Therapeutic opportunities within the DNA damage response. Nat Rev Cancer.

Pink RC, Samuel P, Massa D, Caley DP, Brooks SA, Carter DRF. 2015. The passenger strand, miR-21-3p, plays a role in mediating cisplatin resistance in ovarian cancer cells. Gynecol Oncol 137: 143–151. https://pubmed.ncbi.nlm.nih.gov/25579119/ (Accessed January 19, 2021).

Rice TW, Patil DT, Blackstone EH. 2017. 8th edition AJCC/UICC staging of cancers of the esophagus and esophagogastric junction: Application to clinical practice. Ann Cardiothorac Surg 6: 119–130. https://pubmed.ncbi.nlm.nih.gov/28447000/ (Accessed March 5, 2021).

Richards D, Kocs DM, Spira AI, David McCollum A, Diab S, Hecker LI, Cohn A, Zhan F, Asmar L. 2013. Results of docetaxel plus oxaliplatin (DOCOX) ± cetuximab in patients with metastatic gastric and/or gastroesophageal junction adenocarcinoma: Results of a randomised Phase 2 study. Eur J Cancer 49: 2823–2831. https://pubmed.ncbi.nlm.nih.gov/23747051/ (Accessed January 14, 2021).

Rizvi NA, Hellmann MD, Snyder A, Kvistborg P, Makarov V, Havel JJ, Lee W, Yuan J, Wong P, Ho TS, et al. 2015. Mutational landscape determines sensitivity to PD-1 blockade in non-small cell lung cancer. Science (80-).

Ronellenfitsch U, Schwarzbach M, Hofheinz R, Kienle P, Kieser M, Slanger TE, Jensen K, Burmeister B, Kelsen D, Niedzwiecki D, et al. 2013. Perioperative chemo(radio)therapy versus primary surgery for resectable adenocarcinoma of the stomach, gastroesophageal junction, and lower esophagus. Cochrane Database Syst Rev.

Ruhstaller T, Thuss-Patience P, Hayoz S, Schacher S, Knorrenschild JR, Schnider A, Plasswilm L, Budach W, Eisterer W, Hawle H, et al. 2018. Neoadjuvant chemotherapy followed by chemoradiation and surgery with and without cetuximab in patients with resectable esophageal cancer: A randomized, open-label, phase III trial (SAKK 75/08). Ann Oncol 29: 1386–1393. https://pubmed.ncbi.nlm.nih.gov/29635438/ (Accessed January 14, 2021).

Saunders CT, Wong WSW, Swamy S, Becq J, Murray LJ, Cheetham RK. 2012. Strelka: Accurate somatic small-variant calling from sequenced tumor-normal sample pairs. Bioinformatics 28: 1811–1817. https://pubmed.ncbi.nlm.nih.gov/22581179/ (Accessed February 15, 2021).

Schenck RO, Lakatos E, Gatenbee C, Graham TA, Anderson ARA. 2019. NeoPredPipe: High-throughput neoantigen prediction and recognition potential pipeline. BMC Bioinformatics.

Secrier M, Li X, De Silva N, Eldridge MD, Contino G, Bornschein J, Macrae S, Grehan N, O’Donovan M, Miremadi A, et al. 2016. Mutational signatures in esophageal adenocarcinoma define etiologically distinct subgroups with therapeutic relevance. Nat Genet 48: 1131–1141.

Sharma M. 2015. Ilorasertib in Treating Patients With CDKN2A-deficient Advanced or Metastatic Solid Cancers That Cannot Be Removed by Surgery - Full Text View - ClinicalTrials.gov. https://clinicaltrials.gov/ct2/show/NCT02540876 (Accessed January 19, 2021).

Sjoquist KM, Burmeister BH, Smithers BM, Zalcberg JR, Simes RJ, Barbour A, Gebski V. 2011. Survival after neoadjuvant chemotherapy or chemoradiotherapy for resectable oesophageal carcinoma: An updated meta-analysis. Lancet Oncol.

Smyth EC, Lagergren J, Fitzgerald RC, Lordick F, Shah MA, Lagergren P, Cunningham D. 2017. Oesophageal cancer. Nat Rev Dis Prim 3. https://pubmed.ncbi.nlm.nih.gov/28748917/ (Accessed March 10, 2021).

Snyder A, Makarov V, Merghoub T, Yuan J, Zaretsky JM, Desrichard A, Walsh LA, Postow MA, Wong P, Ho TS, et al. 2014. Genetic Basis for Clinical Response to CTLA-4 Blockade in Melanoma. N Engl J Med.

Subramanian A, Tamayo P, Mootha VK, Mukherjee S, Ebert BL, Gillette MA, Paulovich A, Pomeroy SL, Golub TR, Lander ES, et al. 2005. Gene set enrichment analysis: A knowledge-based approach for interpreting genome-wide expression profiles. Proc Natl Acad Sci U S A 102: 15545–15550. https://pubmed.ncbi.nlm.nih.gov/16199517/ (Accessed March 9, 2021).

Tan C, Qian X, Guan Z, Yang B, Ge Y, Wang F, Cai J. 2016. Potential biomarkers for esophageal cancer. Springerplus 5.

Tao CJ, Lin G, Xu YP, Mao WM. 2015. Predicting the response of neoadjuvant therapy for patients with esophageal carcinoma: An in-depth literature review. J Cancer 6: 1179–1186.

Turkington RC, Knight LA, Blayney JK, Secrier M, Douglas R, Parkes EE, Sutton EK, Stevenson L, McManus D, Halliday S, et al. 2019. Immune activation by DNA damage predicts response to chemotherapy and survival in oesophageal adenocarcinoma. Gut.

Uboveja A, Satija YK, Siraj F, Sharma I, Saluja D. 2020. p73 – NAV3 axis plays a critical role in suppression of colon cancer metastasis. Oncogenesis 9: 12. http://www.nature.com/articles/s41389-020-0193-4 (Accessed September 22, 2020).

Van Allen EM, Miao D, Schilling B, Shukla SA, Blank C, Zimmer L, Sucker A, Hillen U, Foppen MHG, Goldinger SM, et al. 2015. Genomic correlates of response to CTLA-4 blockade in metastatic melanoma. Science (80-).

Van Cutsem E, Köhne C-H, Hitre E, Zaluski J, Chang Chien C-R, Makhson A, D’Haens G, Pintér T, Lim R, Bodoky G, et al. 2009. Cetuximab and Chemotherapy as Initial Treatment for Metastatic Colorectal Cancer. N Engl J Med 360: 1408–1417. https://pubmed.ncbi.nlm.nih.gov/19339720/ (Accessed January 14, 2021).

Van Loo P, Nordgard SH, Lingjærde OC, Russnes HG, Rye IH, Sun W, Weigman VJ, Marynen P, Zetterberg A, Naume B, et al. 2010. Allele-specific copy number analysis of tumors. Proc Natl Acad Sci U S A.

van Velzen MJM, Derks S, van Grieken NCT, Haj Mohammad N, van Laarhoven HWM. 2020. MSI as a predictive factor for treatment outcome of gastroesophageal adenocarcinoma. Cancer Treat Rev 86. https://doi.org/10.1016/j.ctrv.2020.102024 (Accessed January 25, 2021).

Wong C, Law S. 2017. Predictive factors in the evaluation of treatment response to neoadjuvant chemoradiotherapy in patients with advanced esophageal squamous cell cancer. J Thorac Dis.

Yang H, Wang K. 2015. Genomic variant annotation and prioritization with ANNOVAR and wANNOVAR. Nat Protoc 10: 1556–1566. https://pubmed.ncbi.nlm.nih.gov/26379229/ (Accessed January 29, 2021).

Zhao S, Wang Y, Cao L, Ouellette MM, Freeman JW. 2010. Expression of oncogenic K-ras and loss of Smad4 cooperate to induce the expression of EGFR and to promote invasion of immortalized human pancreas ductal cells. Int J Cancer.

Zhu J, Muskhelishvili L, Tong W, Borlak J, Chen M. 2020. Cancer genomics predicts disease relapse and therapeutic response to neoadjuvant chemotherapy of hormone sensitive breast cancers. Sci Rep.

