## Supplementary figures and images for "Genomic analysis of response to neoadjuvant chemotherapy in esophageal adenocarcinoma"

### Supplementary Figure S1

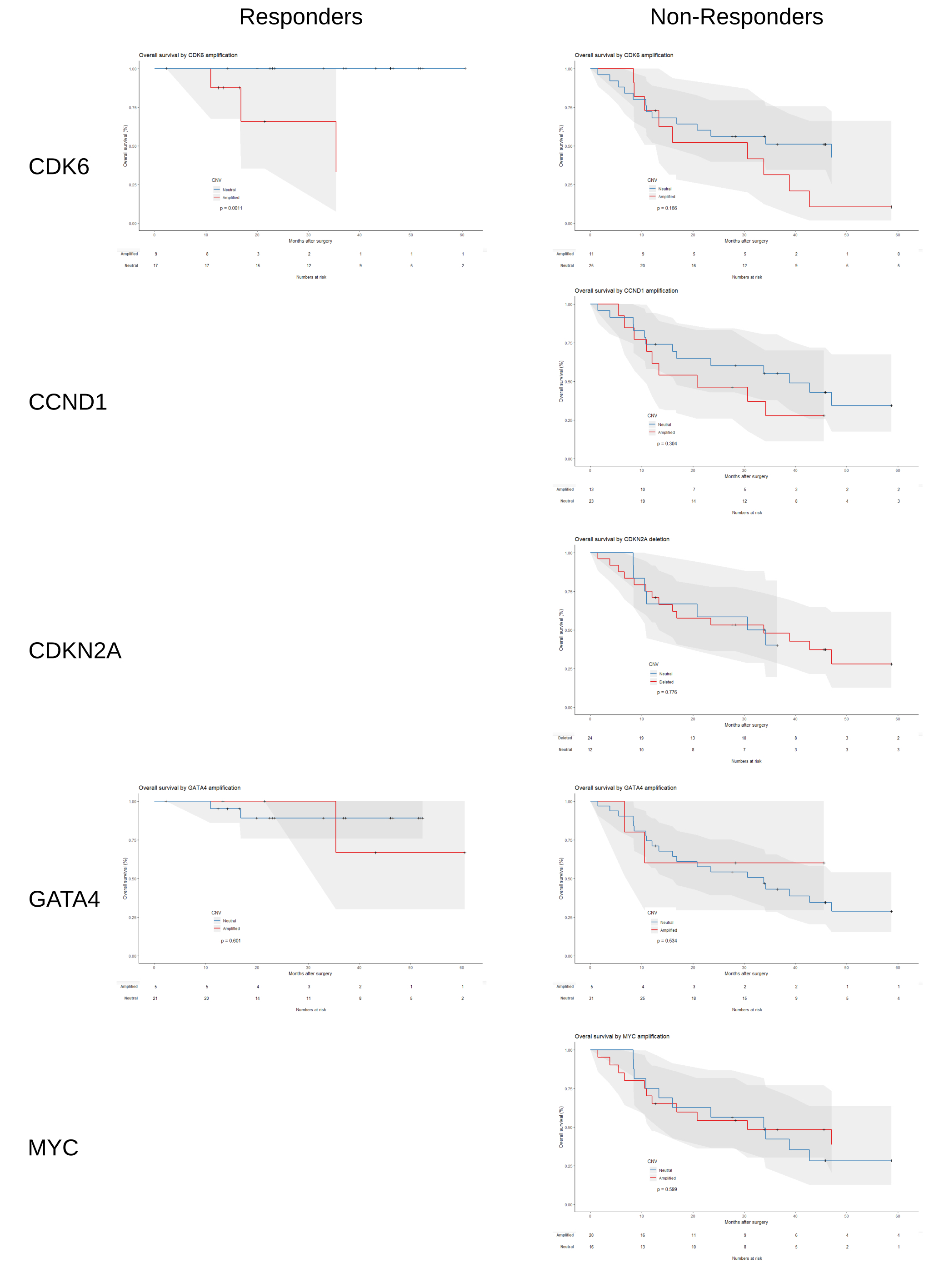
